# Tracing boundaries in Eastern Antarctica: Multi-scale drivers of soil microbial communities across the hyperarid Vestfold Hills

**DOI:** 10.1101/2021.09.22.461446

**Authors:** Eden Zhang, Paul Czechowski, Aleks Terauds, Sin Yin Wong, Devan S. Chelliah, Mark Raven, Mark M. Tanaka, Belinda C. Ferrari

**Author notes:** **Author contributions:** BCF, AT, MMT, DSC and EZ designed the study. AT collated the environmental predictor data and coordinated sample collection. MR conducted XRF analyses. PC developed the analysis code then both EZ and SYW revised it. EZ processed the sequencing data and performed analyses with assistance from SYW. EZ wrote the manuscript with input from BCF, DSC, PC and AT. All authors read, collaborated and approved the final manuscript.

## Abstract

Microorganisms are key to sustaining core ecosystem processes across terrestrial Antarctica but they are rarely considered in conservation frameworks. Whilst greater advocacy has been made towards the inclusion of microbial data in this context, there is still a need for better tools to quantify multispecies responses to environmental change. Here, we extend the scope of Gradient Forest modelling beyond macroorganisms and small datasets to the comprehensive polar soil microbiome encompassing >17, 000 sequence variants for bacteria, micro-eukarya and archaea throughout the hyperarid Vestfold Hills of Eastern Antarctica. Quantification of microbial diversity against 79 physiochemical variables revealed that whilst rank-order importance differed, predictors were broadly consistent between domains, with greatest sharing occurring between bacteria and micro-eukarya. Moisture was identified as the most robust predictor for shaping the regional soil microbiome, with highest compositional turnover or “splits” occurring within the 10 – 12 % moisture content range. Often the most responsive taxa were rarer lineages of bacteria and micro-eukarya with phototrophic and nutrient-cycling capacities such as *Cyanobacteria* (up to 61.81 % predictive capacity), *Chlorophyta* (62.17 %) and *Ochrophyta* (57.81 %). These taxa groups are thus at greater risk of biodiversity loss or gain to projected climate trajectories, which will inevitably disturb current ecosystem dynamics. Better understanding of these threshold tipping points will positively aid conservation efforts across Eastern Antarctica. Furthermore, the successful implementation of an improved Gradient Forest model also presents an exciting opportunity to broaden its use on microbial systems globally.

## Main Text

Changing climate in the Antarctic drives the expansion and greater connectivity of its ice-free habitats, which predominantly consist of soil microbiota (Lee *et al*., 2017). In response to warming, amelioration of environmental conditions such as increased soil moisture and nutrient content is expected to alter current ecosystem dynamics (Chown *et al*., 2015; McGaughran, Laver and Fraser, 2021). However, it is difficult to predict the long-term impact of disturbance on community composition and functionality, especially given the immense diversity of microorganisms and their highly variable responses (Allison & Martiny, 2008; Cavicchioli *et al*., 2019; Robinson *et al*., 2018 and 2020). This is further exacerbated by a general lack of available studies, especially in Eastern Antarctica (Zhang *et al*., 2020). The hyperarid Vestfold Hills comprise of low-lying hilly landscapes, built upon a mixture of unconsolidated rock and unsorted debris left behind by the retreating East Antarctic Ice Sheet (Hirvas, Nenonen and Quilty, 1993). Strewn throughout its glacially scoured basins are numerous bodies of water, which have been the primary focus for biodiversity, evolutionary and ecological studies (Laybourn-Parry and Bell, 2014; Wilkins *et al*., 2013). In contrast, its soil microbiome remains poorly investigated (Zhang *et al*., 2020). Moving beyond descriptive reports, better understanding on the relationship between soil microbial diversity and the edaphic environment will be crucial towards achieving sound conservation planning and management across terrestrial Antarctica (Bergstrom *et al*., 2020; Groves *et al*., 2002; Rath *et al*., 2018).

Soil micro-niches are highly variable and complex (Fierer, 2017). Shifts in microbial community composition are expected to occur at extremely fine scales (Chong *et al*., 2015; Cowan *et al*., 2014; Ferrari *et al*., 2016; Siciliano *et al*., 2014). Great interest therefore lies in quantifying key drivers of diversity and threshold tipping points along edaphic gradients, which enables the formation of a solid basis for predicting shifts in response to complex environmental change (Kennicutt II *et al*., 2014; Rath *et al*., 2018). This knowledge may then be used to critically inform the designation of protected areas in order to effectively manage Antarctica’s unique biodiversity (Lee *et al*., 2017). Inclusion of microorganisms, however, has been rare in the development of Antarctica conservation frameworks as well as other ecosystems around the globe (Cavicchioli *et al*., 2019; Hughes *et al*., 2015 and 2018). As such, there is an increasingly urgent need to find or develop suitable methods to bridge this conservation gap – one option being Gradient Forest (Ellis, Smith and Pitcher, 2012).

Based on an aggregation of Random Forest (Breiman, 2001), which only considers the response of single species, Gradient Forest offers a flexible approach to explore multispecies responses between biodiversity and the environment (Ellis, Smith and Pitcher, 2012). Accounting for complex interactions by fitting multiple regression models, Gradient Forest quantifies the relative importance of different predictors and their non-linear correspondence with community compositional turnover or “splits” along predictor gradients, which can be upscaled across extensive geographic areas to define spatial groups that capture variation in species composition and turnover (Stephenson *et al*., 2018). This output may be used to critically inform conservation planning and management, especially where biodiversity survey data is sparse (Pitcher *et al*., 2012). However, the application of such modelling approaches has thus far been limited to studies on macroorganisms in forest or marine ecosystems (Maguire *et al*., 2015; Pitcher *et al*., 2012; Stephenson *et al*., 2018).

For the first time, we employ a modified Gradient Forest framework on the polar soil microbiome (i.e., bacteria, micro-eukarya and archaea), using reanalysed 16S and 18S rRNA gene amplicon sequence variants (ASVs) obtained from four sites (Adam’s Flat “AF”, Heidemann Valley “HV”, Old Wallow “OW” and Rookery Lake “RL”) throughout the hyperarid Vestfold Hills of Eastern Antarctica (Zhang *et al*., 2020). Doing so, we (1) quantify the relative importance of 79 measured physiochemical predictors on the local and regional distribution of polar soil microbial communities across the region; and (2) identify where any major compositional shifts occur along important predictor gradients in order to define spatial boundaries and abiotic thresholds. These outcomes will be used to prioritize sensitive areas for further sampling and targeted microbial studies across the Vestfold Hills, with significant global outreach to other arid soil environments.

## Results

### Soil characteristics and phyla are distributed unevenly between and within sites along spatially explicit transects

Overlap of samples on the environmental nMDS indicate that measured soil physiochemical variables are generally conserved across the Vestfold Hills (Figure 1). However, significant (ANOSIM *p* <0.05) local heterogeneity is still apparent, for example elevation (*R* = 0.81), aspect (*R* = 0.45) and moisture (*R* = 0.27) amongst other nutrient-related factors (Table 1). In terms of biodiversity, bubbleplots depicting the relative distribution and abundance of bacteria, micro-eukarya and archaea revealed that a handful of phyla dominate the regional soil microbiome (Figures 2 and S1). These include members belonging to the ubiquitous and metabolically diverse *Actinobacteria, Bacteroidetes, Proteobacteria* and *Chloroflexi* as well as a relatively high abundance (>60 %) of unclassified micro-eukarya (Figure S2a and b). Rarer phyla (<10 %) accounted for the main phylogenetic difference amongst sites (Figures 2 and S1), which include an uneven distribution of *Acidobacteria* (start of AF transect), *Cyanobacteria* (start of RL transect), *Ca. Eremiobacterota* (Start of HV transect) and micro-algae such as *Ochrophyta* (mid of OW transect) and *Chlorophyta* (start of RL transect). In contrast, archaea were generally consistent between and within sites (Figures 2c and S1c). Overall, greatest phylogenetic diversity was observed at OW (100/102 m) and RL (0/2 m) (Figure 2).

**Figure 1.**
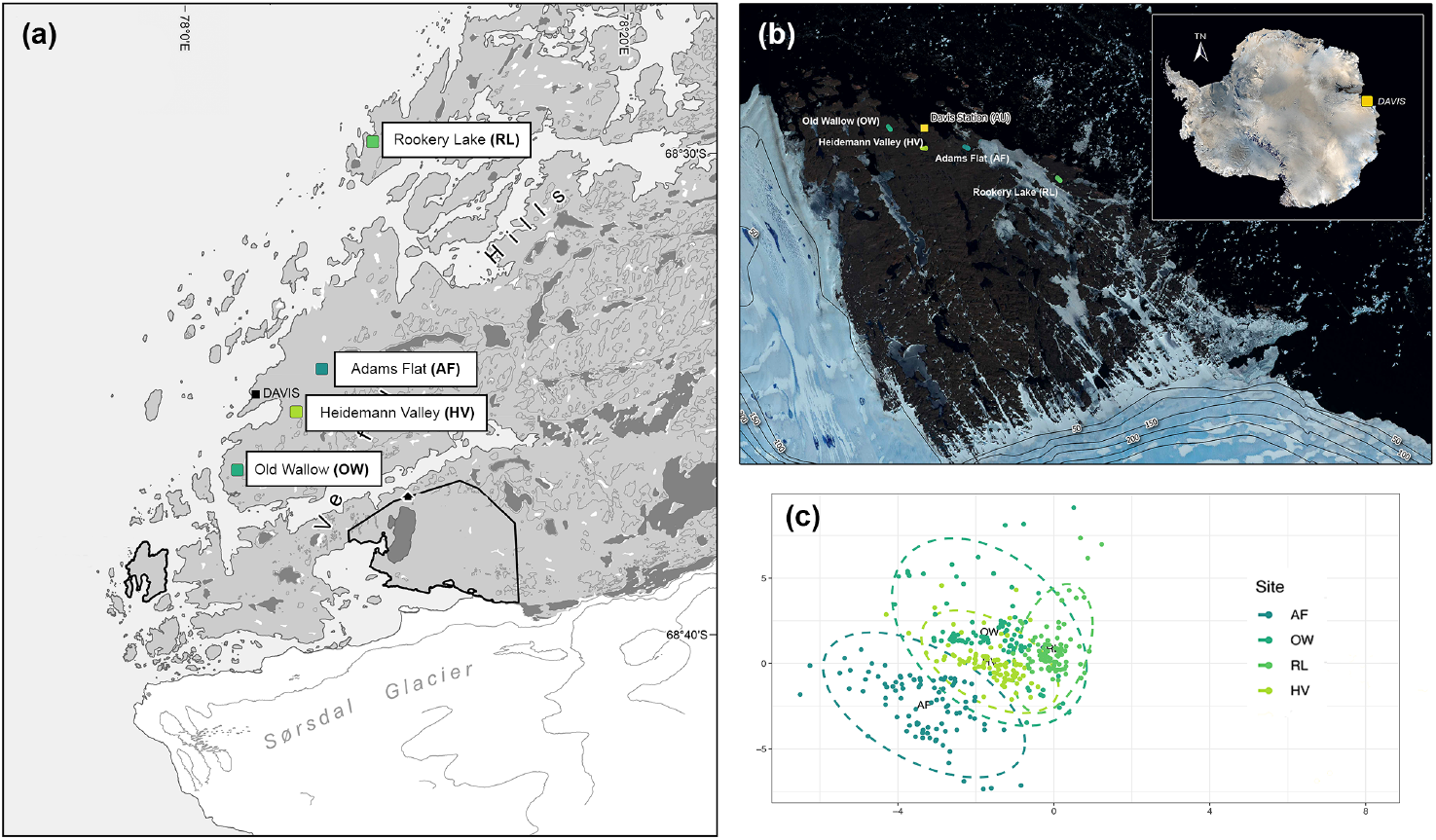
Overview of the four sampling sites (AF, HV, OW and RL) throughout the hyperarid Vestfold Hills in Eastern Antarctica as presented by (a) map (AAD map catalogue No. 14, 399), (b) satellite image (Quantarctica v3 GIS package in QGIS 3.4.7) and (c) environmental nMDS. Measured soil physiochemical variables (79) are generally conserved across the region with visible overlap amongst all sites, especially between OW, HV and RL whereas greater variability is observed for samples at AF.

**Figure 2.**
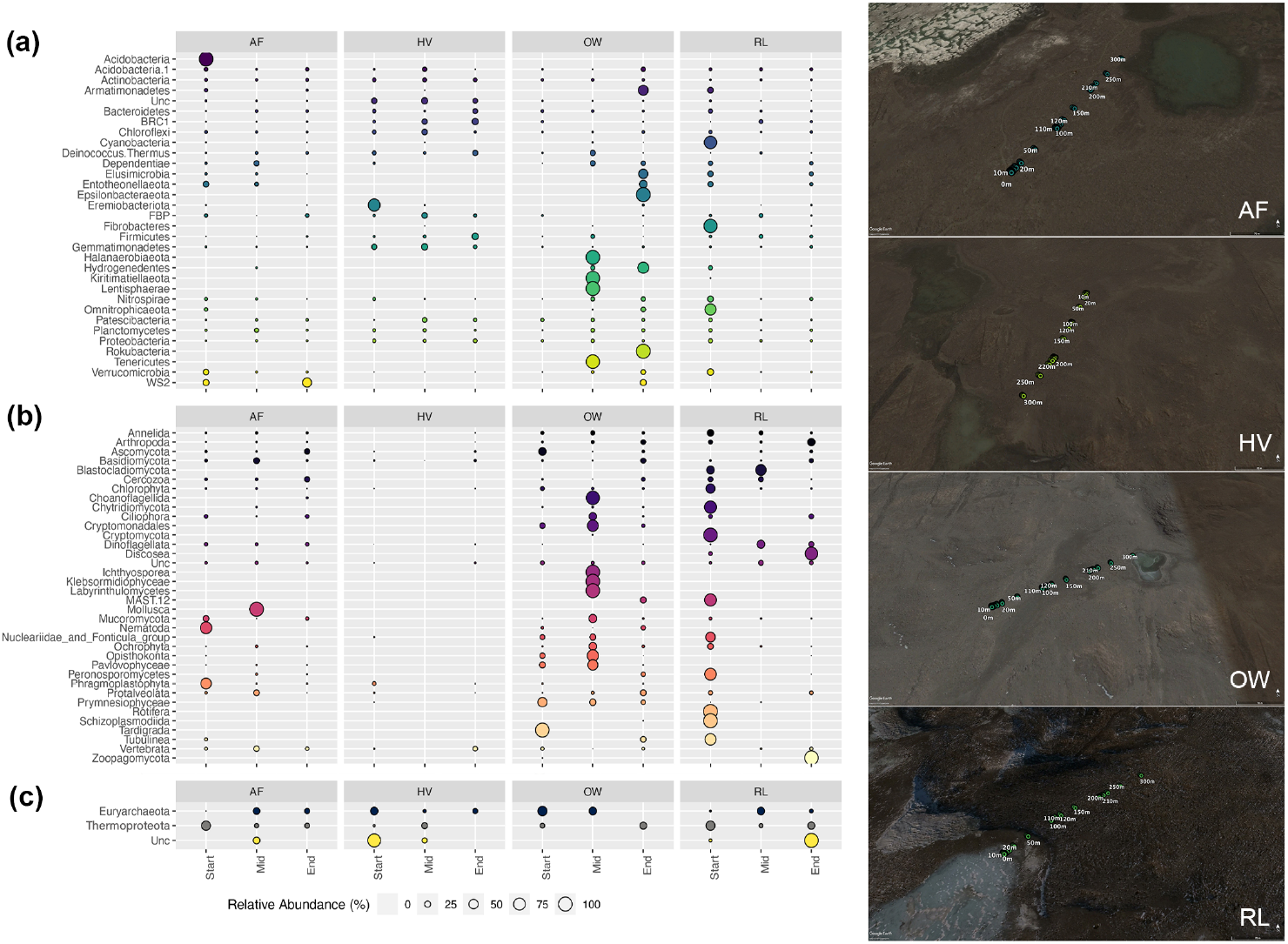
Relative distribution of each phylum across the Vestfold Hills region for polar soil (a) bacteria, (b) micro-eukarya and (c) archaea, where start (0/2 m), mid (100/102 m) and end (200/202 m) indicate the position of samples along transects at each site. Phyla are unevenly distributed between and within sites along their respective transects, with greatest phylogenetic diversity observed at OW (mid) and RL (start).

**Table 1.**
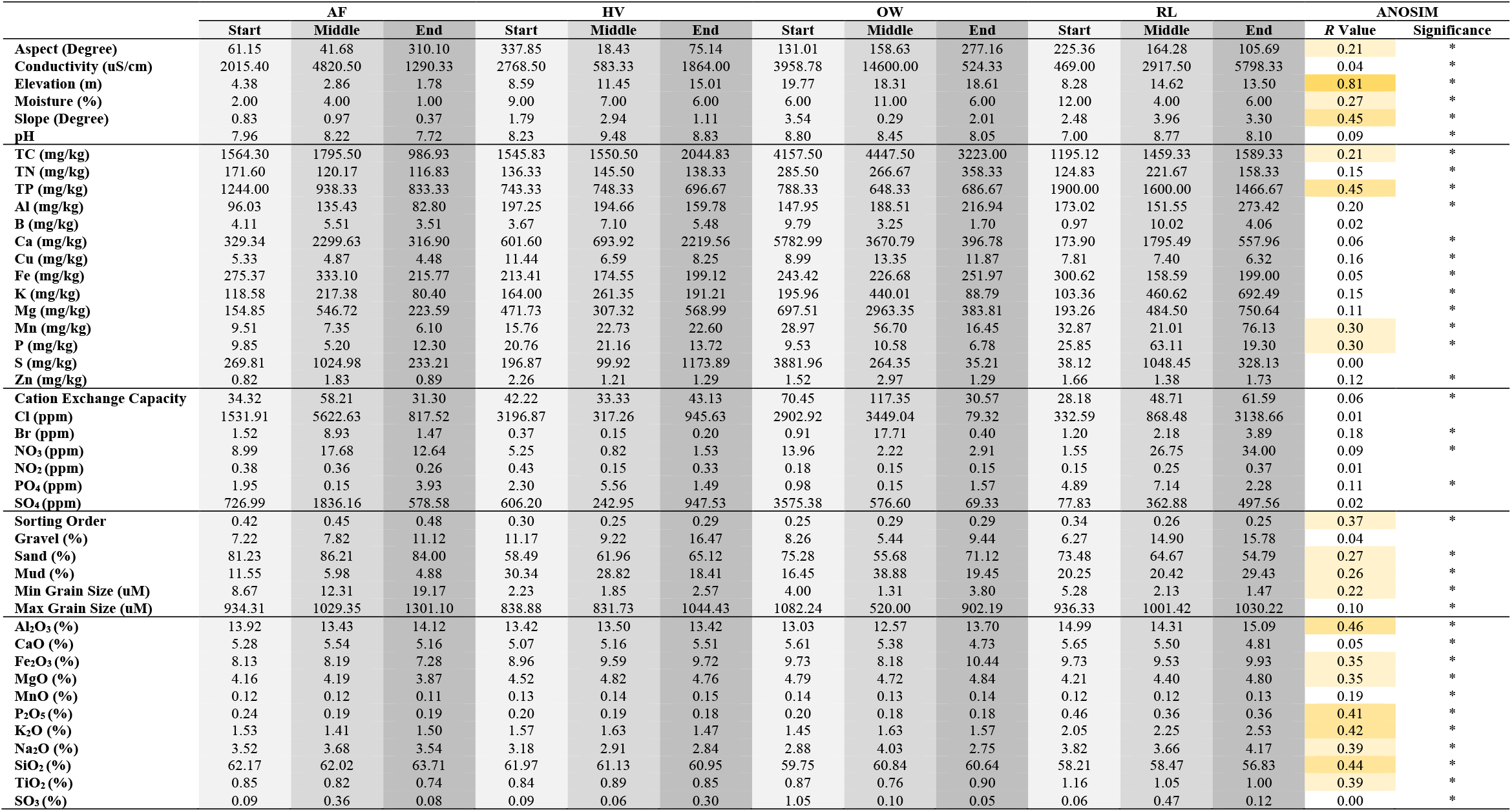
Summary of complete environmental predictors averaged (n = 18) between sites. Where start (0/2 m), mid (100/102 m) and end (200/202 m) indicate position of samples (n = 6) along the transects and (*) denotes a significant (*p* <0.05) difference for that variable between sites based on ANOSIM using 9999 permutations.

### Moisture availability shapes the hyperarid Vestfold Hills soil microbiome

Overall importance plots generated from Gradient Forest analyses demonstrate that rank-order importance differed but predictors were generally consistent between domains (Figure 3). Moisture was identified as the most robust predictor in shaping the regional soil microbiome, followed by nutrient-related factors (e.g., Br and K), pH and aspect amongst other predictors of intermediate importance (Figure 3). On the local scale (Table 2), differences between sites reflect local physiochemical variations (Table 1). Some examples include calcium (Ca) at AF (Figure S2), phosphate-related factors (e.g., PO_4_, P and TP) at HV (Figure S3), conductivity at OW (Figure S4) and boron (B) content at RL (Figure S5). In general, aspect and trace elements (e.g., B, Zinc “Zn”, iron “Fe” and manganese “Mn”) became more important as differentiating factors for local communities. Bacteria and micro-eukarya generally shared a similar set of predictors whereas archaea remained largely distinct, except at OW and RL (Figure 3 and Table 2). Overall, the suite of environmental variables predicted a relatively modest fraction of variation in the distribution of polar soil bacteria (*R*^2^ = 0.009 – 0.732), micro-eukarya (*R*^2^ = 0.005 – 0.622) and archaea (*R*^2^ = 0.031 – 0.300) across the Vestfold Hills (Table S1). Of which these variables had some predictive capacity (i.e., positive *R*^2^) for phyla on both regional (bacteria = 27, micro-eukarya = 21, archaea = 2) and local (bacteria = 4 – 21, micro-eukarya = 2 – 12, archaea = 1 – 3) scales (Tables S1 and S2).

**Figure 3.**
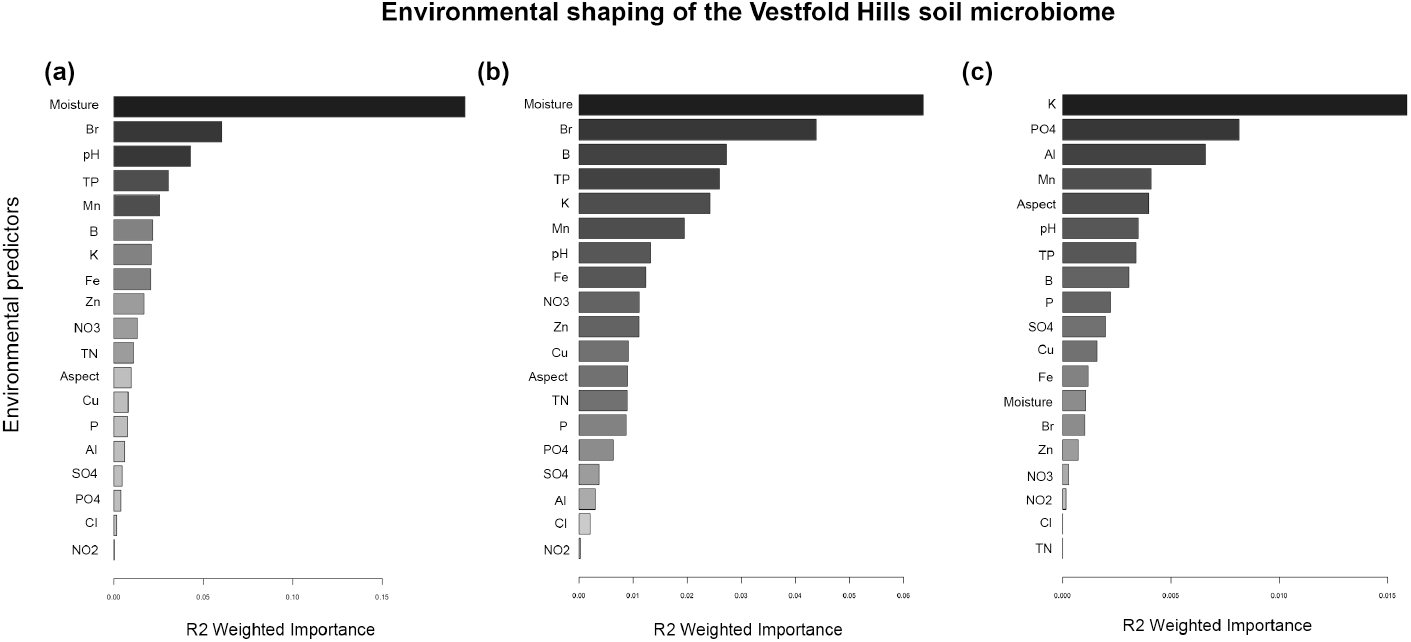
Overall importance (R^2^ weighted) of predictors driving regional distributions of polar soil (a) bacteria, (b) micro-eukarya and (c) archaea across the Vestfold Hills. The suite of environmental variables predicted a moderate fraction of variation in regional species distribution. Although rank-order importance differed, most predictors were relatively consistent across the three domains, especially between bacteria and micro-eukarya. Soil moisture and nutrient-related factors (e.g., Br and K) along with pH were identified as the most robust predictors of regional diversity.

**Table 2.**
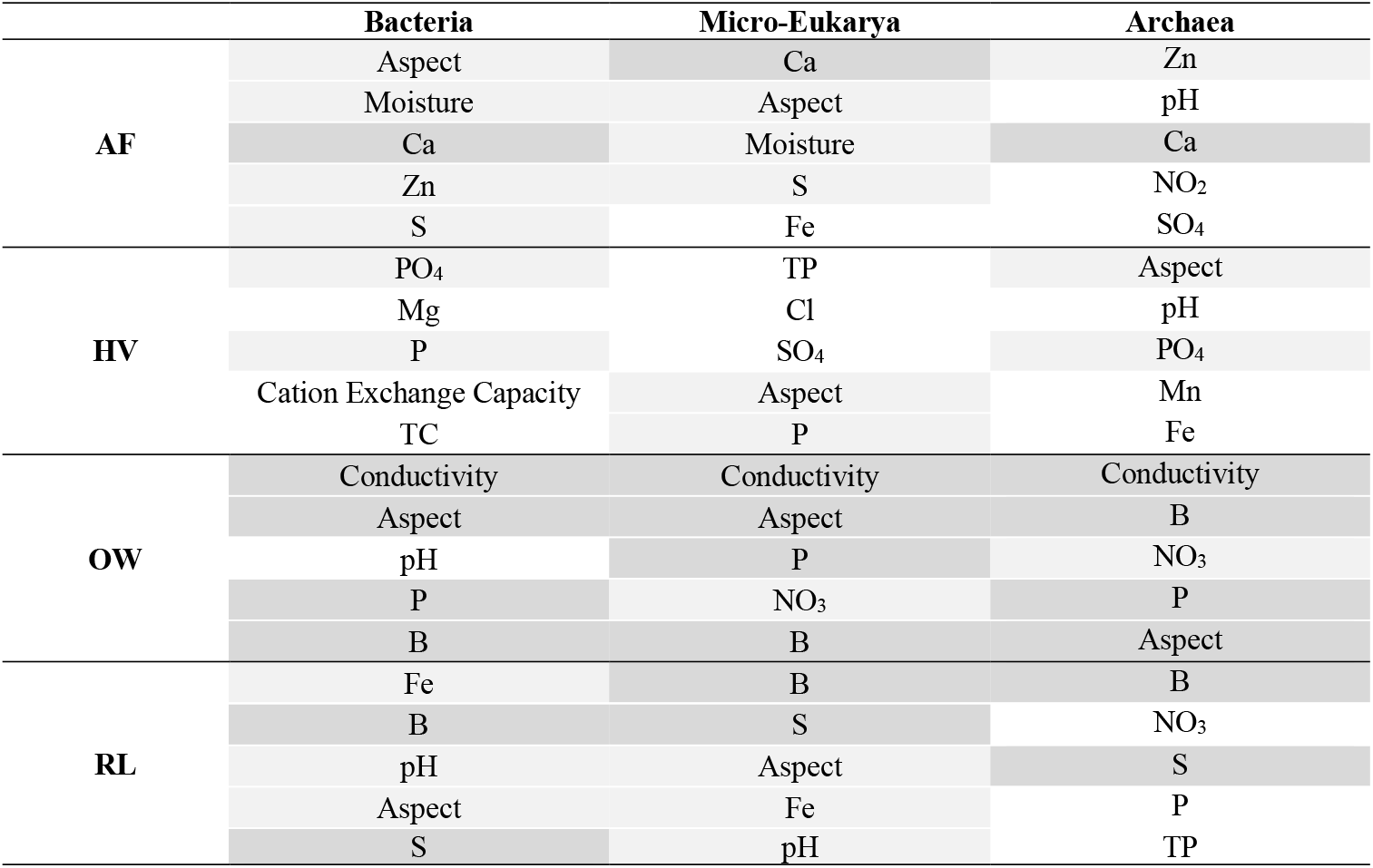
Top five most robust predictors for local distributions of polar soil bacteria, micro-eukarya and archaea. Where shaded boxes indicate sharing of predictor(s) across two or more domains.

|  | **Bacteria** | **Micro-Eukarya** | **Archaea** |
| --- | --- | --- | --- |
| **AF** | Aspect | Ca | Zn |
|  | Moisture | Aspect | pH |
|  | Ca | Moisture | Ca |
|  | Zn | S | NO<sub>2</sub> |
|  | S | Fe | SO<sub>4</sub> |
| **HV** | PO<sub>4</sub> | TP | Aspect |
|  | Mg | Cl | pH |
|  | P | SO<sub>4</sub> | PO<sub>4</sub> |
|  | Cation Exchange Capacity | Aspect | Mn |
|  | TC | P | Fe |
| **OW** | Conductivity | Conductivity | Conductivity |
|  | Aspect | Aspect | B |
|  | pH | P | NO<sub>3</sub> |
|  | P | NO<sub>3</sub> | P |
|  | B | B | Aspect |
| **RL** | Fe | B | B |
|  | B | S | NO<sub>3</sub> |
|  | pH | Aspect | S |
|  | Aspect | Fe | P |
|  | S | pH | TP |

### Gradient responses and thresholds vary between domains and over different spatial scales

Cumulative importance plots reveal that changes in community composition or “splits” along edaphic gradients were non-uniform and responses largely varied between phyla over both regional and local scales (Figures 4 and S2 – 5). For example, along the moisture gradient, many important splits occurred in the range *c*. 10 – 12 % (Figure 4), indicating that high rates of compositional turnover (e.g., *Cyanobacteria, Ochrophyta* and *Chlorophyta*) corresponded with small changes in moisture content across the region (Figure 4a and b). All sites sampled throughout the Vestfold Hills are extremely arid, with AF being the driest (2.00 % ± 2.00) followed by HV (7.00 % ± 2.00) whilst soils at OW (8.00 % ± 4.00) and RL (8.00 % ± 4.00) retain higher moisture content (Table 1). Higher abundances of *Cyanobacteria, Ochrophyta* and *Chlorophyta* were detected at OW and RL (Figures 2a/b and S1a/b). Similarly, along the Br gradient, many important splits occurred in the range *c*. 5 – 15 ppm affecting regional abundances of rarer phyla like *Kiritimatiellaeota* and *Ichthyosporea* (Figure 4a and b), which were primarily distributed towards the middle (17.71 Br ppm) of the OW transect (Figure 2a and b). Regarding archaea, particularly *Thermoproteota* (Figures 2c and S1c), its regional distribution and abundance did not change much along the K gradient despite exhibiting a steep response *c*. 600 – 700 ppm (Figure 4c).

**Figure 4.**
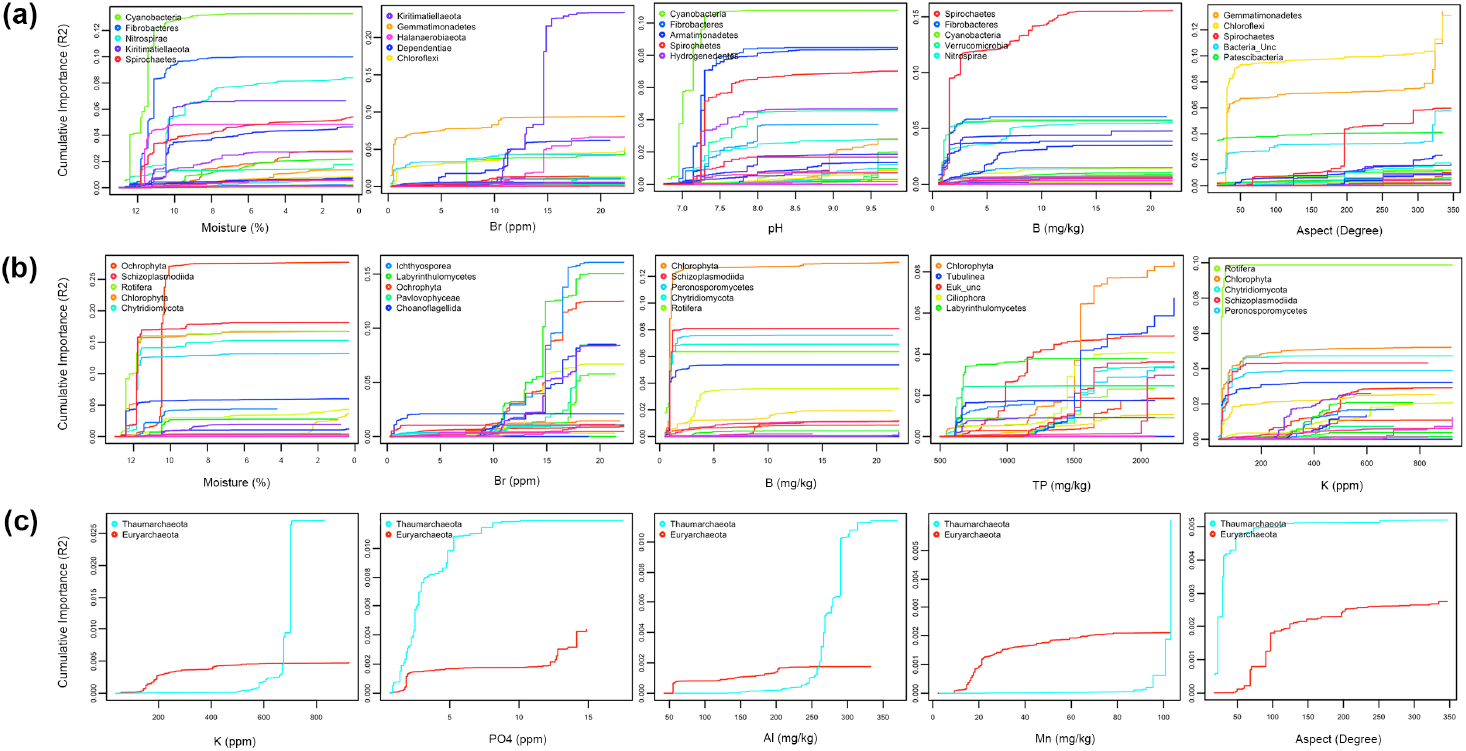
Cumulative plots visualising species turnover or “splits” along gradients of the top five most robust predictors for the regional distribution of polar soil (a) bacteria, (b) micro-eukarya and (c) archaea across the Vestfold Hills, where each line denotes a separate phylum and steeper slopes indicate higher rates of composition turnover. Splits along environments gradients were generally non-uniform, indicating variables rates of change in species composition. More pronounced transitions were observed for bacteria (e.g., *Kiritimatiellaeota, Cyanobacteria, Spirochetes* and *Gemmatimonadetes*) and micro-eukarya (e.g., *Ochrophyta, Ichthyosporea, Chlorophyta* and *Rotifera*). For bacteria, notable splits occurred between the following ranges: *c*. 5–15 bromide “Br” (ppm), *c*. 10–12 % moisture, *c*. 7.0–7.5 pH, *c*. 0–5 boron “B” (mg/kg) and at 50/200^0^ aspect (degree). For micro-eukarya, notable splits occurred between the following ranges: *c*. 10–12 % moisture, *c*. 10–15 Br ppm, *c*. 2 B mg/kg, *c*. 1500–1600 total phosphorous “TP” (mg/kg) and *c*. 50 potassium “K” (ppm). For archaea, notable splits occurred between the following ranges: *c*. 600–700 K ppm, *c*. 0–5 phosphate “PO_4_” (ppm), *c*. 250 aluminium “Al” (mg/kg), *c*. 100 manganese “Mn” (mg/kg) and c. 0/100^0^ aspect.

In comparison, notable differences in compositional responses were observed at finer scales (Figures S2 – 5). For instance, along the Ca gradient at AF, important splits occurred in the range *c*. 100–1200 mg/kg (Figure S2), affecting *FBP* (*c*. 0 – 200 Ca mg/kg) at steeper rates than *Labyrinthulomycetes, Ochrophyta* and unclassified archaea (*c*. 1000 – 1200 Ca mg/kg), where higher abundances of *FBP* are distributed towards the start (329.34 Ca mg/kg) and end (316.90 Ca mg/kg) of the AF transect (Figure 2a). Whilst at HV, along the PO_4_ gradient, important splits occurred in the range *c*. 2 – 8 ppm (Figure S3), mainly affecting *Chloroflexi* (c. 2 – 6 PO_4_ ppm) and *Euryarchaeota* (c. 6 – 8 PO_4_ ppm), with *Chloroflexi* present in higher abundances towards the middle (5.56 PO_4_ ppm) of the HV transect (Figure 2a and c). Similarly, along the conductivity gradient at OW, important splits occurred in the range *c*. 0 – 15, 000 uS/cm (Figure S4), mainly affecting *Deinococcus-Thermus* (*c*. 7500 – 10, 000 uS/cm), *Ochrophyta* (*c*. 10, 000 – 15, 000 uS/cm) and *Euryarchaeota* (*c*. 2500 – 5000 uS/cm), where higher abundances of all three phyla were distributed towards the middle (14, 600 uS/cm) of the OW transect (Figure 2). At RL, important splits along the B gradient occurred in the range *c*. 0 – 12 mg/kg (Figure S5), affecting *Cyanobacteria* (*c*. 2 – 4 B mg/kg) and *Chlorophyta* (*c*. 0 – 2 B mg/kg) at steeper rates than *Euryarchaeota* (*c*. 6 – 10 B mg/kg), where higher abundances of *Cyanobacteria* and *Chlorophyta* were detected towards the start (0.97 B mg/kg) of the RL transect.

### Classification of spatial groups for regional conservation planning

Mapping of expected biodiversity patterns based on information about gradient responses and thresholds previously stored in cumulative importance curves identified an average of two-to-four spatial groups on both regional and local scales (Figures 5 and S6 – 8). For the most part, the Vestfold Hills soil microbiome appeared to be a fairly homogenous environment though greater heterogeneity in compositional response and overall phylogenetic diversity was detected at OW and RL (Figures 2 and 5). In particular, their soil bacterial and micro-eukaryotic components situated towards the start (RL; Figure 5b) and middle (OW; Figure 5a and b) of the transects where retainment of soil moisture (11 – 12 %) is highest in the region (Table 1). On finer scales (Figures S6 – 8), biological responses were notably more heterogenous with visible boundaries between the start, middle and end of transects – with aspect playing a key role. However, when taking into consideration sample size and variation, predictive power remains overall more useful on the broad regional scale for identifying priority areas for further studies.

**Figure 5.**
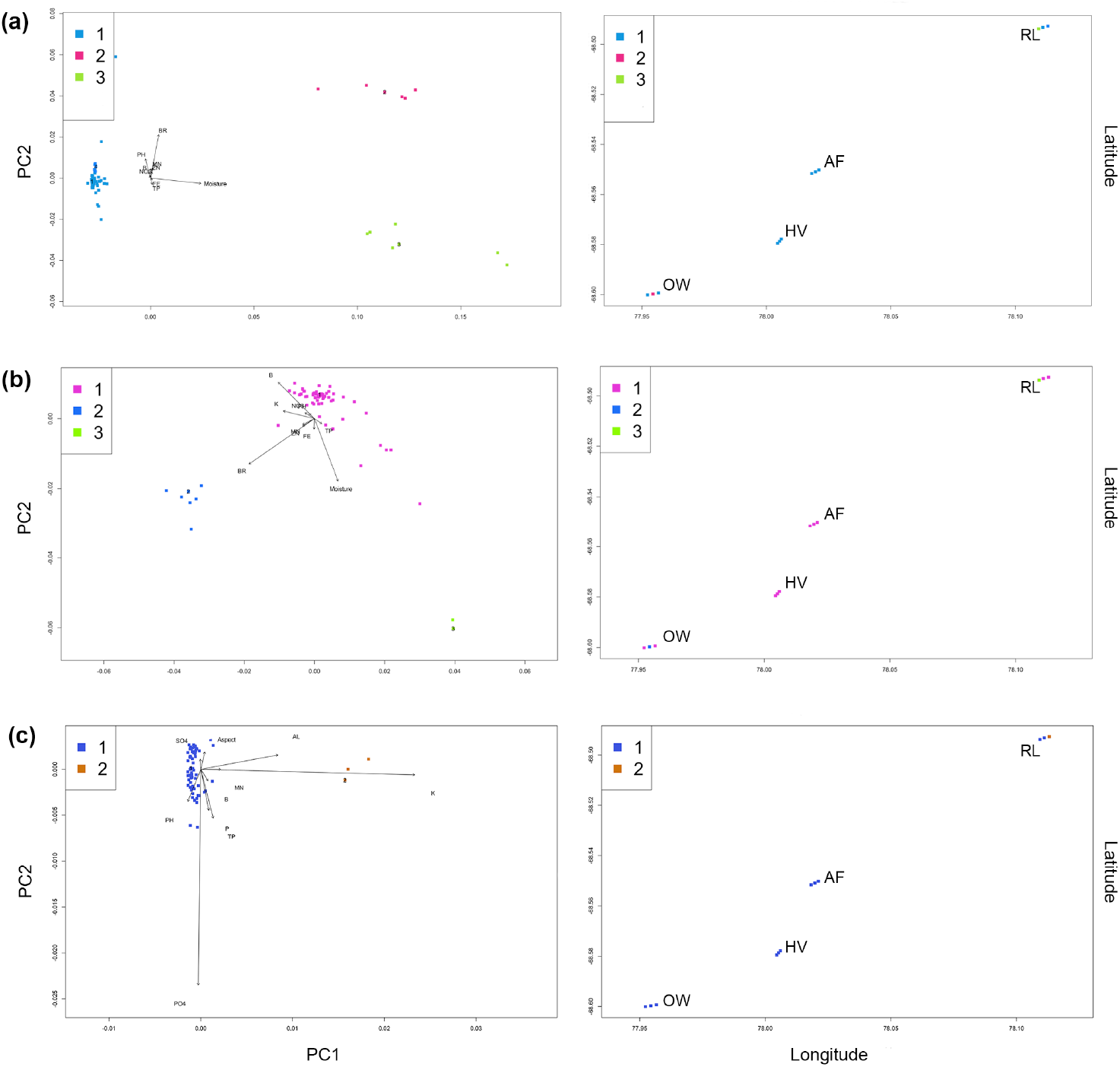
Biological and geographical bi-plots classifying spatial groups for polar soil (a) bacteria (3), (b) micro-eukarya (3) and (c) archaea (2) across the Vestfold Hills (n = 72), where different colours capture variations in species composition and turnover. Spatial patterns of turnover in soil biodiversity across the region described by cluster classification are strongly related to moisture content and nutrient-related factors, except for archaea. Greater heterogeneity in response groups were generally observed at OW (mid of transect for bacteria and micro-eukarya) and RL (start of transect for micro-eukarya and end of transect for archaea).

## Discussion

Robust discrimination of key drivers underlying biogeographic patterns of polar soil microbial communities offered by Gradient Forest make it well suited as primary input towards systematic conservation planning across the hyperarid Vestfold Hills. Distinctive gradient responses and thresholds for bacterial and micro-eukaryotic communities at Old Wallow (OW) and Rookery Lake (RL) has led to the prioritisation of these two East Antarctic sites as conservation targets for further sampling (Figures 2, 4 and 5). More in-depth studies, however, are needed to critically inform potential management actions. Nonetheless, our findings provide a baseline for anticipating shifts in microbial response to ameliorated conditions as consequence of regional warming (Robinson *et al*., 2020; McGaughran, Laver and Fraser, 2021).

Environmental shaping of the hyperarid Vestfold Hills soil microbiome is primarily driven by variances in moisture throughout the region (Figure 3), which is largely regulated by extrinsic factors such as temperature (Delgado-Baquerizo *et al*. 2018). Along the regional moisture gradient (Figure 4), strongest rates of compositional turnover were observed for rarer lineages of bacteria (e.g., *Cyanobacteria*) and micro-eukarya (e.g., *Chlorophyta* and *Ochrophyta*) with phototrophic and/or nutrient-cycling capacities such as nitrogen and hydrogen metabolism (Bothe *et al*., 2010; Fernandes *et al*., 2016; Fuerst & Sagulenko 2012; Lea-Smith *et al*., 2016). These taxa groups were found in relatively high abundances at OW and RL (Figures 2 and S1), likely due to the presence of meltwater lakes and stochastic nutrient inputs from wildlife (Huang & Zhu, 2009). Projected warming trends may favour the rapid growth of photosynthetic potential across terrestrial Antarctica (Bay, Greening and Ferrari, 2018). An example being in the form of microalgal blooms near bird or seal colonies, like those at OW and RL, which may also act as important terrestrial carbon sinks (Gray *et al*., 2020). Cascading effects from this hypothetical scenario would inevitably disturb current ecosystem dynamics, especially when taking into consideration that <0.1 % of the frigid landmass is vegetated (Fretwell *et al*., 2011). Therefore, placing extreme selective pressure against chemoautrophic lineages (e.g., *Actinobacteria, Ca. Eremiobacterota* and *Ca. Dormibacterota*) and thereby leading to potential competition for dominance between primary production strategies across Eastern Antarctica (Bay *et al*., 2021; Bay *et al*., 2021; Ji *et al*., 2017; Ray *et al*., 2020).

Ecosystem resilience, however, remains a poorly understood concept due to the lack of empirical data (Allison & Martiny, 2008). As such, it is unknown whether compositional changes will also affect functionality nor the extent to which it drives biodiversity gain or loss over larger spatial scales, especially within the context of a warming Antarctica (Lee et al., 2017). Outputs from Gradient Forest analyses provided some predictive capacity, up to 73.2 % in some cases for bacteria and micro-eukarya depending on spatial scale (Table S1), but offered limited resolution on archaea across the hyperarid Vestfold Hills. This suggests that the environment may not be the primary driver of distributions and compositions for some taxa. As such, spatial planning and management activities also need to consider these other processes that drive biodiversity patterns (Pitcher *et al*., 2012). However, further sampling and individual site-level studies are necessary to draw more explicit conclusions – by starting at OW and RL.

Building upon previous work highlighting the unique microbial biodiversity found in Eastern Antarctica (Zhang *et al*., 2020), we advance methods to better apprehend their key edaphic drivers and threshold tipping points on multiple scales. Increased capacity of Gradient Forest to model strain-level data enables the integration of diverse microbial communities in Antarctic conservation and management actions, including the development of new protected areas (Hughes, Cowan and Wilmotte, 2015), thus taking a crucial step towards achieving sound conservation and management across terrestrial Antarctica (Cavicchioli *et al*., 2019; Hughes, Cowan and Wilmotte, 2015). Moreover, our modified Gradient Forest framework will also be valuable for assessing disturbance, rehabilitation and restoration within other terrestrial environments around the globe.

## Methods

### Study area and sampling design

Sampling in the Vestfold Hills region (centred 68°55”S, 78°25”E) was performed in 2012 by expeditioners via the Australian Antarctic Program (AAP) across four locations within vicinity of the Australian Davis research station (68°35”S, 77°58”E) (Figure 1). These include: Adams Flat (AF: 68°33”S, 78°1”E); Heidemann Valley (HV: 68°35”S, 78°0”E); Old Wallow (OW: 68°36”S, 77°58”E) and Rookery Lake (RL: 68°36”S, 77°57”E). At each site, samples (n = 18) were taken from the top 10 cm of soil along three parallel transects at the following distance points of 0, 2, 100, 102, 200 and 202 m (Siciliano *et al*., 2014). For the purpose of this study, samples were not pooled but categorised into start (0/2 m), mid (100/102 m) and end (200/202 m) according to their location along the transect.

### Soil physiochemical analysis

All soils (n = 72) included in this study had been submitted by the Australian Antarctic Division (AAD) for extensive physiochemical analysis using standard procedures as performed by Bioplatforms Australia (BPA). As described in Siciliano *et al*. (2014; Appendix A. Supplementary Data) and Table S3, this includes the following: pH, conductivity, moisture, aspect, elevation, slope, calcium “Ca”, magnesium “Mg”, potassium “K”, sulfur “S”, total carbon “TC”, total nitrogen “TN”, total phosphorous “TP”, aluminium “Al”, boron “B”, copper “Cu”, iron “Fe”, manganese “Mn”, zinc “Zn”, cation exchange capacity, chloride “Cl”, bromide “Br”, nitrate “NO_3_”, nitrite “NO_2_”, phosphate “PO_4_”, phosphorous “PO_3_”, sulphate “SO_4_”, sorting order, gravel, sand, mud, min grain size, max grain size, aluminium oxide “Al_2_O_3_”, calcium oxide “CaO”, iron (III) oxide “Fe_2_O_3_”, magnesium oxide “MgO”, manganese oxide “MnO”, phosphorous pentoxide “P_2_O_5_”, potassium oxide “K_2_O”, sodium oxide “Na_2_O”, silicon oxide “SiO_2_”, sulfur trioxide “SO_3_” and titanium oxide “TiO_2_”.

### DNA extraction and Illumina amplicon sequencing

Soil samples were extracted in triplicate using the FASTDNA™ SPIN Kit for Soil (MP Biomedicals, Santa Ana, CA, US) and Qubit™ 4 Fluorometer (ThermoFisher Scientific, NSW, Australia) as per manufacturers” instructions. Diluted DNA (1:10 using nuclease-free water) was submitted to the Ramaciotti Centre for Genomics (UNSW Sydney, Australia) for amplicon paired-end sequencing on the Illumina MiSeq platform (Illumina, California, US) with controls in accordance to protocols from the AusMicrobiome project (Bissett *et al*., 2016). For bacteria and archaea, we respectively targeted the V1–3 hypervariable region of 16S rRNA gene sequences using the 27F/519R (Lane, 1991) and A2F/519R primer sets (Reysenbach *et al*., 1995). For micro-eukarya, we targeted the V9 hypervariable region of 18S rRNA gene sequences using the 1391f /EukBr primer set (Amaral-Zettler *et al*., 2009).

### Reprocessing of amplicon sequencing data

Forward and reverse primer sequences were removed from the raw fastq files using cutadapt v2.10 (Martin, 2011). The trimmed paired end reads were merged using FLASH 2 (Magoč & Salzberg, 2011) then converted into fasta format using SEQTK (https://github.com/lh3/seqt). Individual files were concatenated and imported into MOTHUR (Schloss et al., 2009) to screen the sequences, remove ambiguous bases and those with homopolymer runs >8 bp. UPARSE/UNOISE2 (Edgar, 2013 and 2016) was then used dereplicate and denoise the screened sequences, yielding a set of amplicon sequence variants (ASVs – Callahan *et al*., 2017) and abundance count tables. For both 16S and 18S rRNA amplicon gene datasets, taxonomy was classified against the SILVA database release 132 (Quast *et al*., 2013).

### Summary of input data for Gradient Forest analysis

In total, we recovered 51, 978, 909 bacterial 16S rRNA, 17, 994, 697 micro-eukaryotic 18S rRNA and 5, 150, 065 archaeal 16S rRNA gene sequences from the high-throughput amplicon sequencing of the Vestfold Hills soil microbiome (n = 72). After read-quality filtering, a total of 51, 026, 375 bacterial sequences remained, which clustered into 12, 658 ASVs (Table S2). In terms of micro-eukarya, a total of 9, 630, 123 sequences remained after read-quality filtering, which clustered into 1, 507 ASVs (Table S2). Meanwhile, a total of 2, 159, 440 high-quality archaeal sequences were left and these clustered into 367 ASVs (Table S2). Good overall coverage was obtained for bacteria (average reads per sample = 75, 811) micro-eukarya (average reads per sample = 52, 785) and archaea (average reads per sample = 69, 703). All environmental parameters (79) were checked for normality and no transformations were applied, though high-correlated variables (*R*^2^ >0.7) or those with incomplete data were manually removed, resulting in 43 complete predictors (Table S3).

### The R environment

All multivariate and statistical analyses were carried out in R Studio version 4.0.3 (R Core Team, 2020). Unless specified otherwise, all plots were visualised using a combination of ggplot2 v3.1.0 (Wickham, 2016) and ggpubr v0.2 (Kassambara, 2018).

### Gradient Forest analysis

Datasets were stored as individual phyloseq objects (McMurdie & Holmes, 2013). These were prepared by normalising the abundance data using DESeq2 (Love, Huber and Anders, 2014), then agglomerating taxa at the phylum level and extracting defined metadata variables as individual list elements for binary conversion and/or the removal of highly co-correlated variables (*R*^2^ >0.7) using the cor function in R. The R package gradientForest (Ellis *et al*., 2010) was called upon to fit the model (rp1 = 1000, corr.threshold = 0.5) to our optimised list object and return a gradientForest object, which was used to generate a series of plots: (1) predictor overall importance plots (shows mean accuracy importance and the mean importance weighted by species *R*^2^); (2) species cumulative plots (shows cumulative change in abundance of individual species, where changes occur on the gradient and the species changing most on each gradient) and; (3) bi-plots (shows transformed predictors that have been classified into spatial groups, which capture variation in species composition and turnover mapped onto biological and geographical space).

### Deposition of data in an open-source database

All input files including associated environmental metadata and relevant code have been uploaded onto GitHub and is publicly available via https://github.com/Eden3416110/gradientForest.git.

## Competing interests

The authors declare that they have no competing interests.

## ACKNOWLEDGEMENTS

The authors would like to thank the Australian Antarctic Program and all expeditioners for the collection of soil samples (supported by Australian Antarctic Projects 3338 and 5097). We acknowledge funding support through Belinda Ferrari’s Australian Research Council Future Fellowship (FT170100341), the Australian Antarctic Science Grants scheme (AAS 4406 – Ferrari, AAS 4296 – Terauds), and Ausmicrobiome for the provision of the Vestfold Hills biodiversity data.

## Notes

https://github.com/Eden3416110/gradientForest.git

### Competing Interest Statement

The authors have declared no competing interest.

