## Supplementary Information for "Tracing boundaries in Eastern Antarctica: Multi-scale drivers of soil microbial communities across the hyperarid Vestfold Hills"

**This file contains**

Figure S1 to S8

Tables S1 to S2

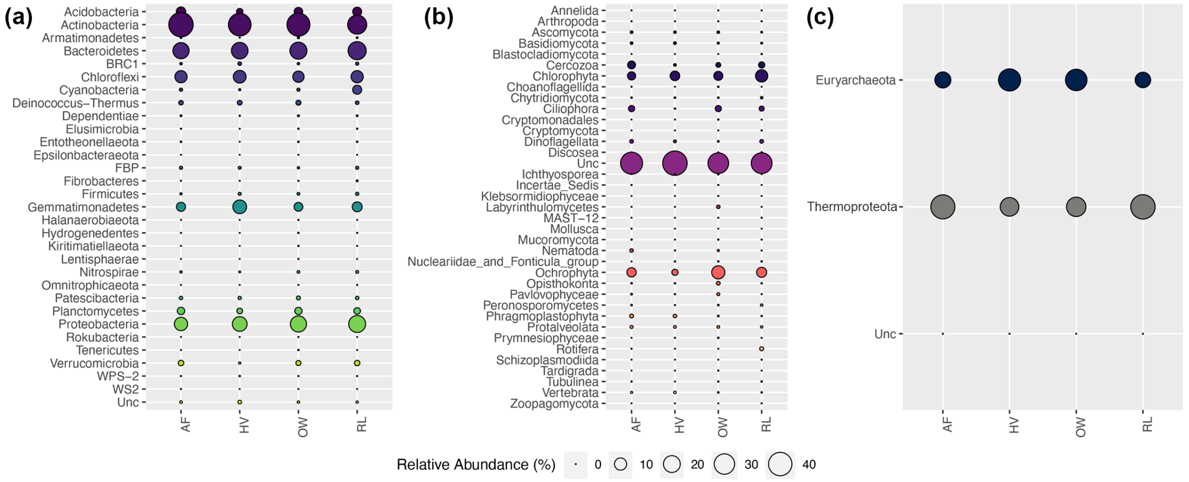


**Figure S1. Relative abundance of phyla between sites for polar soil (a) bacteria, (b) micro-eukarya and (c) archaea across the Vestfold Hills.** A handful of phyla dominates the regional soil microbiome and rarer phyla (<1 %) account for the main phylogenetic difference amongst sites.

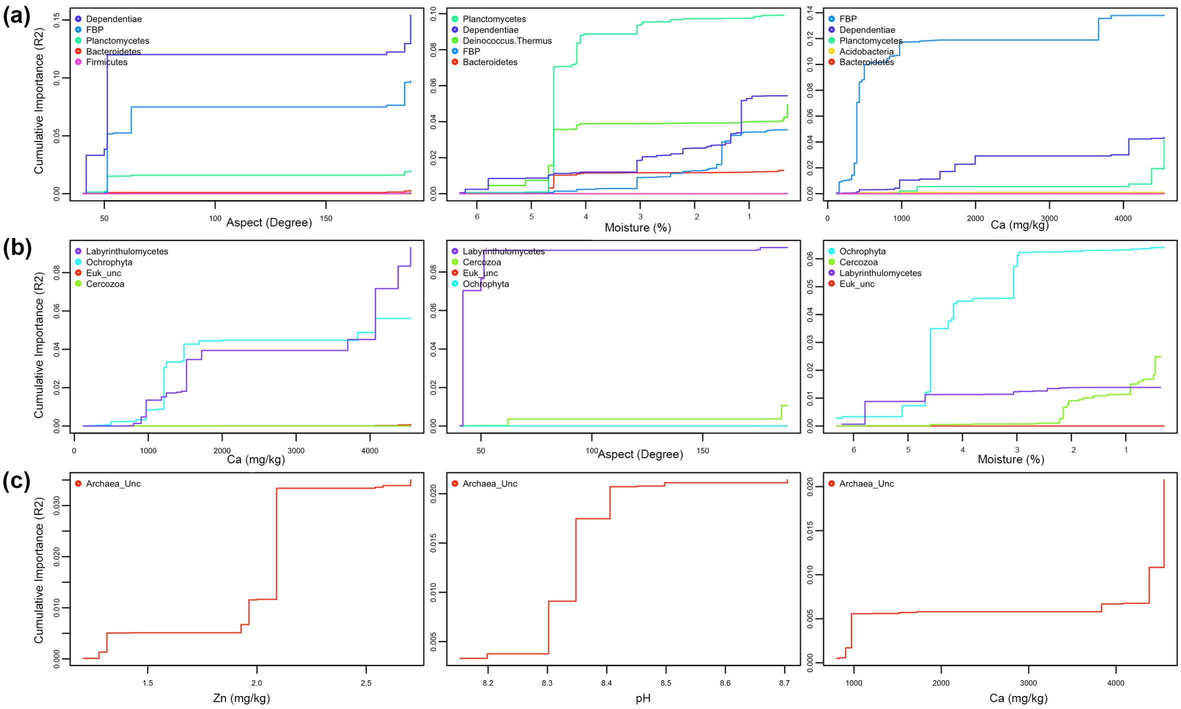


**Figure S2. Cumulative plots for the top five most robust predictors of (a) bacterial, (b) micro-eukaryotic and (c) archaeal communities at Adam's Flat (AF), where each line denotes a separate phylum and steeper slopes indicate higher rates of composition turnover.** Bacterial communities at AF are primarily driven by aspect, moisture content and calcium (Ca), the most responsive phyla include *Dependentiae*, *FBP* and *Planctomycetes* across these gradients. Micro-eukarya at the site, namely *Labyrinthulomycetes* and *Ochrophyta*, also responded to a similar set of predictors. Alongside zinc (Zn) and soil pH, Ca was also an important predictor for local archaeal communities, predominantly affecting unclassified phyla.

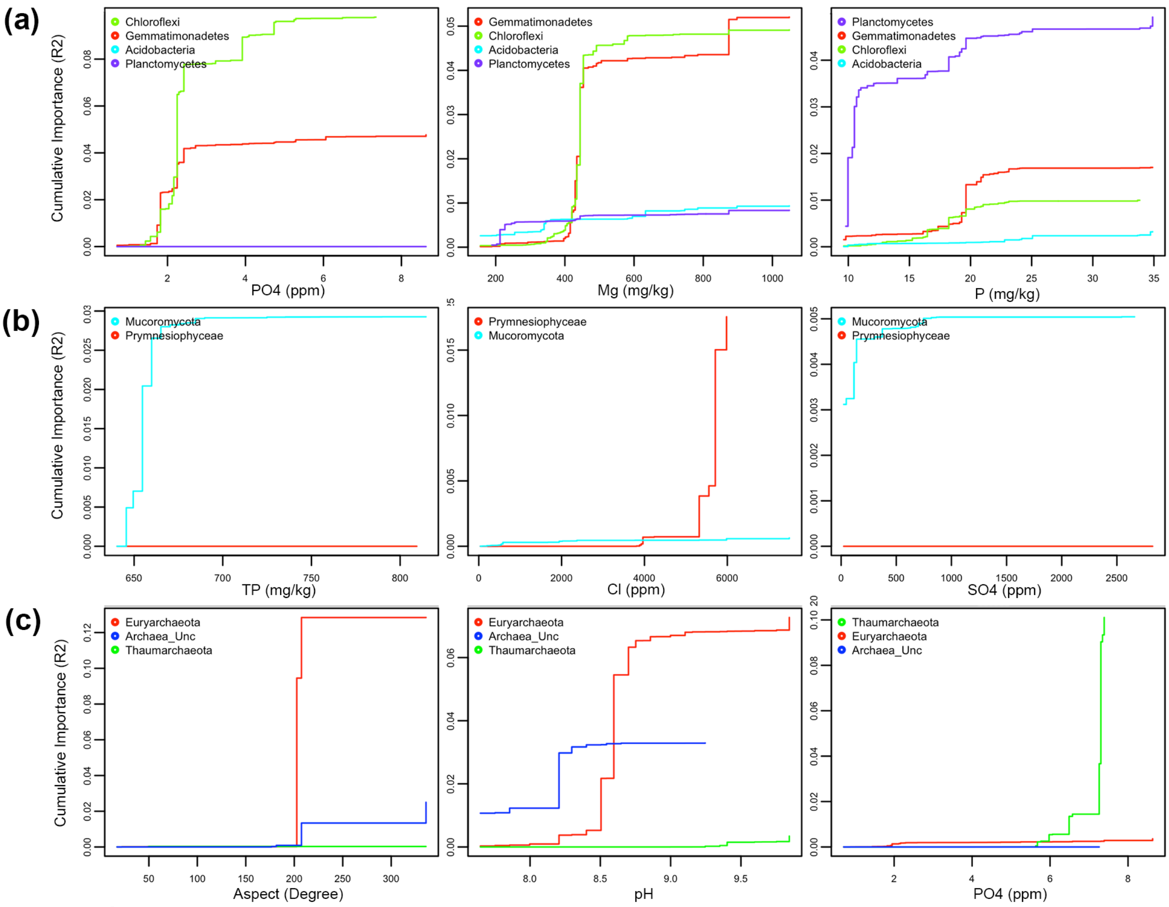


**Figure S3. Cumulative plots for the top five most robust predictors of (a) bacterial, (b) micro-eukaryotic and (c) archaeal communities at Heidemann Valley (HV), where each line denotes a separate phylum and steeper slopes indicate higher rates of composition turnover.** Bacterial communities at HV are primarily driven by phosphate ($PO_4$), magnesium (Mg) and phosphorous (P) content, with phyla such as *Chloroflexi*, *Gemmatimonadetes* and *Planctomycetes* demonstrating the highest compositional turnover rates. Whereas, for micro-eukarya, predictors like total phosphorous (TP), chloride (Cl) and sulphate ($SO_4$) were the most dominant predictors for *Mucoromycota* and *Prymnesiophyceae*. $PO_4$ was also an important predictor for archaea at HV, namely *Thermoproteota*, whilst the greatest change in abundances of *Euryarchaeota* and unclassified phyla were observed along the soil pH gradient and in response to aspect.

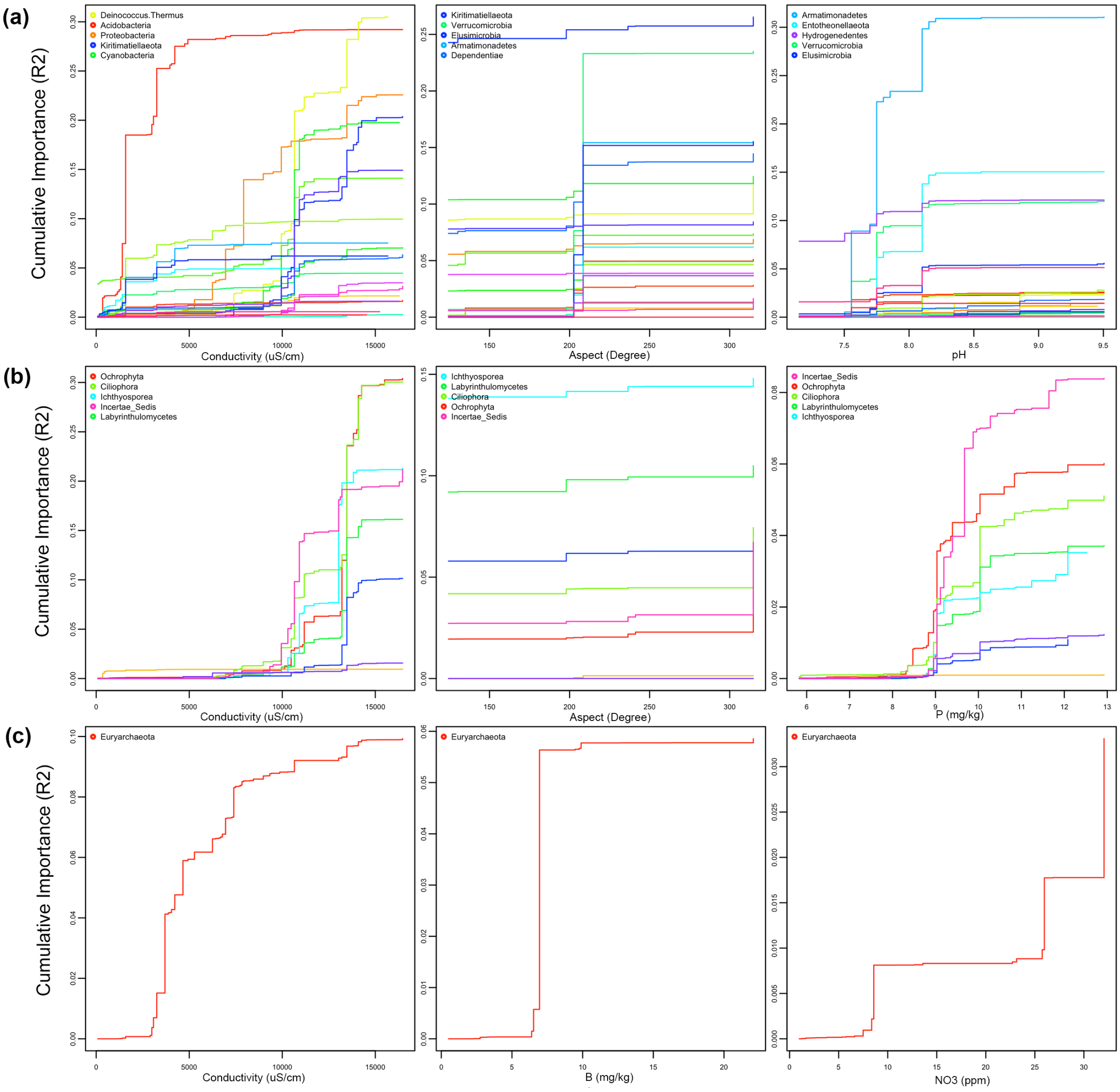


**Figure S4. Cumulative plots for the top five most robust predictors of (a) bacterial, (b) micro-eukaryotic and (c) archaeal communities at Old Wallow (OW), where each line denotes a separate phylum and steeper slopes indicate higher rates of composition turnover.** Bacterial communities at OW are primarily driven conductivity, aspect and soil pH, with *Acidobacteria*, *Kiritimatiellaeota* and *Verrucomicrobia* exhibiting the greatest turnover rates along these gradients. In addition to phosphorous (P), micro-eukarya at the site also responded to a similar set of predictors, mainly affecting abundances of *Ochrophyta* and *Ciliophora* amongst other phyla. Alongside boron (B) and nitrate ($NO_3$), conductivity was also important for local archaeal communities, primarily affecting *Euryarchaeota*.

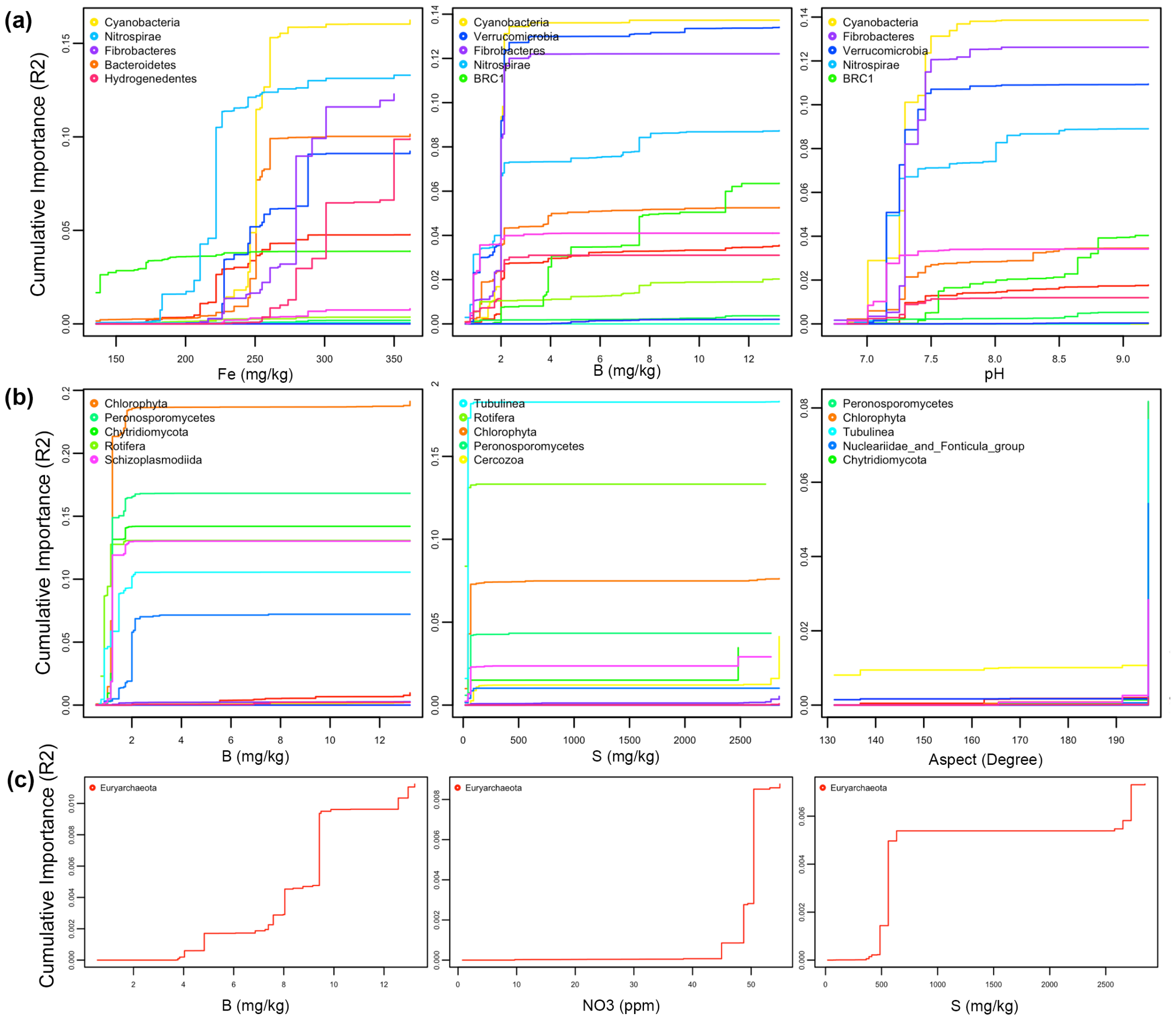


**Figure S5. Cumulative plots for the top five most robust predictors of (a) bacterial, (b) micro-eukaryotic and (a) archaeal communities at Rookery Lake (RL), where each line denotes a separate phylum and steeper slopes indicate higher rates of composition turnover.** Bacterial communities at RL are primarily driven by iron (Fe), boron (B) and soil pH, with *Cyanobacteria*, *Fibrobacteres* and *Nitrospirae* primarily responding to changes along these gradients. Alongside sulphur (S), nitrate ($NO_3$) and aspect, B content was also important for both micro-eukarya and archaea, with *Chlorophyta*, *Peronosporomycetes* and *Euryarchaeota* demonstrating the highest rates of compositional turnover.

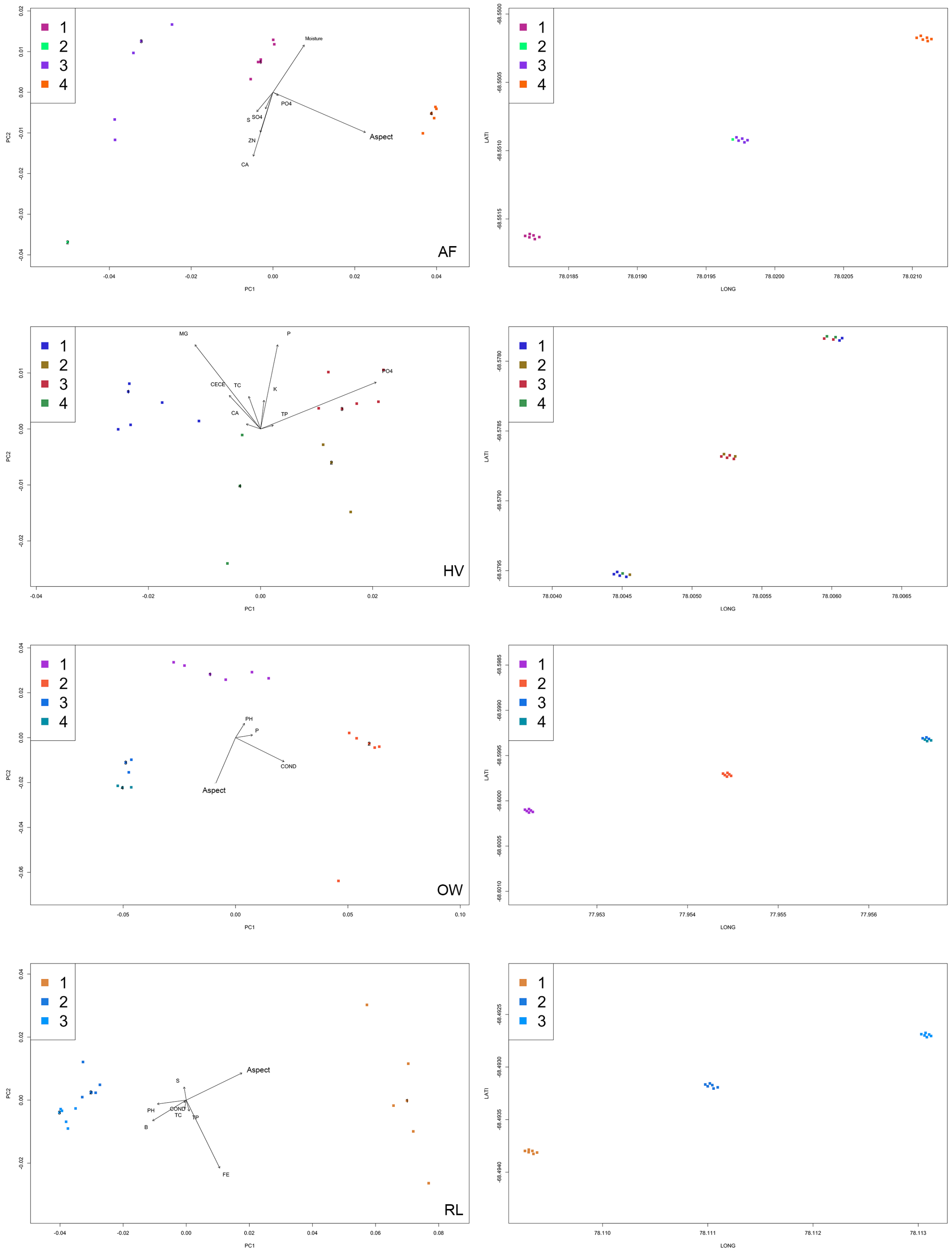


**Figure S6. Biological and geographical bi-plots classifying spatial groups for bacterial communities between sites (n = 18), where different colours capture variations in species composition and turnover.** On average, 3 – 4 spatial groups of bacteria were identified at each site, which were driven by a distinct set of predictors reflecting differences amongst local physiochemical conditions.

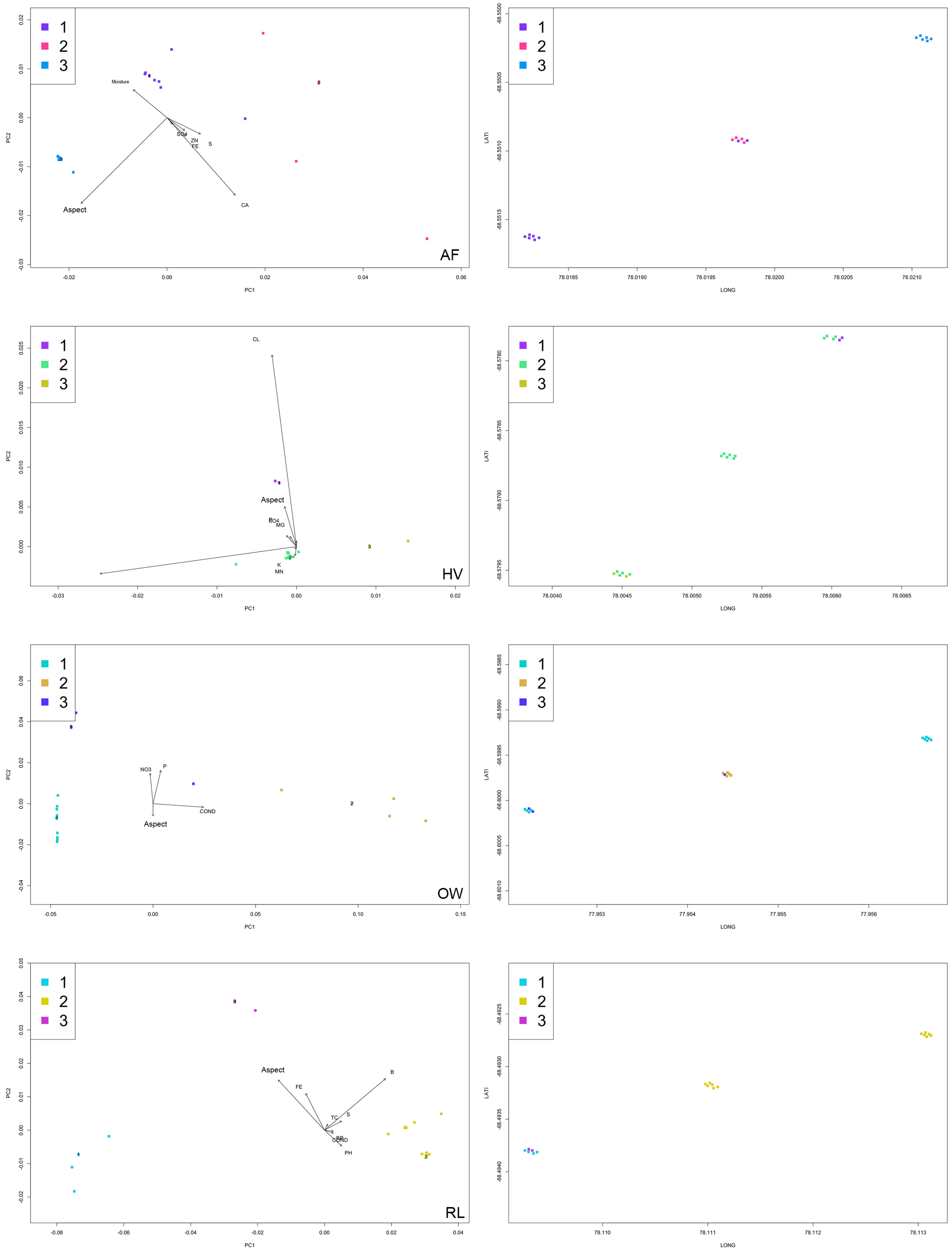


**Figure S7. Biological and geographical bi-plots classifying spatial groups for micro-eukaryotic communities between sites (n = 18), where different colours capture variations in species composition and turnover.** On average, 3 spatial groups of micro-eukarya were identified at each site, which were driven by a distinct set of predictors reflecting differences amongst local physiochemical conditions.

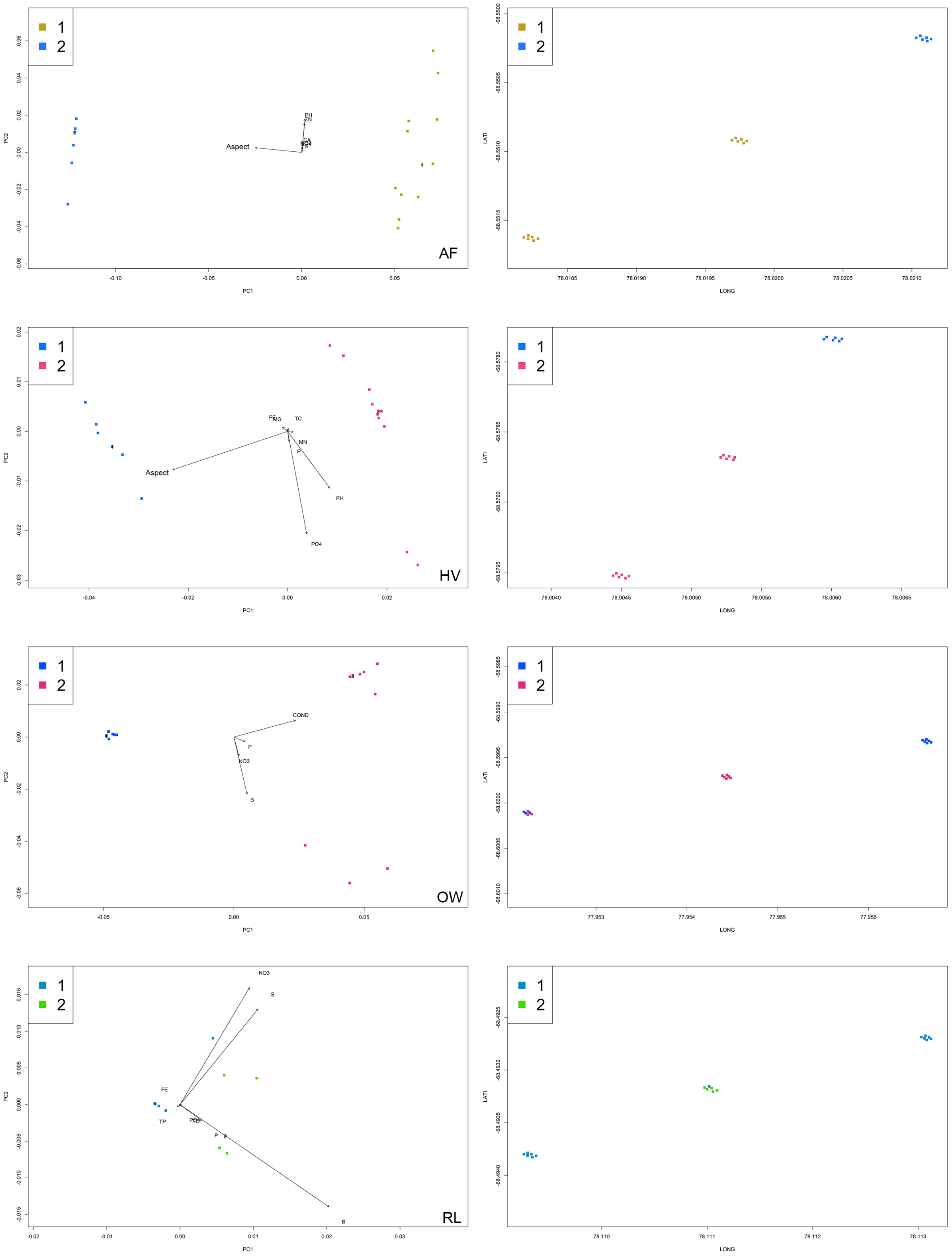


**Figure S8. Biological and geographical bi-plots classifying spatial groups for archaeal communities between sites (n = 18), where different colours capture variations in species composition and turnover.** On average, 2 spatial groups of archaea were identified at each site, which were driven by a distinct set of predictors reflecting differences amongst local physiochemical conditions.

**Table S1. Summary of $R^2$ fitted values at the phylum level.**

| | | $R^2$ Positive | $R^2$ Range (Average) | Top 3 Most Responsive Species |
|---|---|---|---|---|
| Bacteria | **VH** | 27 of 32 phyla | 0.0289–0.592 (0.310) | *Kiritimatiellaeota*<br>*Spirochaetes*<br>*Fibrobacteres* |
| | **AF** | 7 of 28 phyla | 0.0090–0.535 (0.272) | *FBP*<br>*Planctomycetes*<br>*Dependentiae* |
| | **HV** | 4 of 22 phyla | 0.0791–0.240 (0.160) | *Chloroflexi*<br>*Gemmatimonadetes*<br>*Planctomycetes* |
| | **OW** | 21 of 31 phyla | 0.1360–0.732 (0.434) | *Acidobacteria*<br>*Verrucomicrobia*<br>*Gemmatimonadetes* |
| | **RL** | 14 of 27 phyla | 0.0397–0.618 (0.405) | *Cyanobacteria*<br>*Verrucomicrobia*<br>*Nitrospirae* |
| Micro-Eukarya | **VH** | 21 of 38 phyla | 0.005–0.590 (0.298) | *Chlorophyta*<br>*Ochrophyta*<br>*Rotifera* |
| | **AF** | 4 of 33 phyla | 0.022–0.312 (0.167) | *Labyrinthulomycetes*<br>*Ochrophyta*<br>*Cercozoa* |
| | **HV** | 2 of 32 phyla | 0.018–0.048 (0.033) | *Mucoromycota*<br>*Prymnesiophyceae* |
| | **OW** | 8 of 37 phyla | 0.044–0.594 (0.319) | *Ciliophora*<br>*Ochrophyta*<br>*Ichthyosporea* |
| | **RL** | 12 of 34 phyla | 0.013–0.622 (0.318) | *Chlorophyta*<br>*Tubulinea*<br>*Peronosporomycetes* |
| Archaea | **VH** | 2 of 3 phyla | 0.040–0.077 (0.059) | *Euryarchaeota*<br>*Thermoproteota* |
| | **AF** | 1 of 3 phyla | 0.112 | *Unclassified* |
| | **HV** | 3 of 3 phyla | 0.102–0.229 (0.171) | *Euryarchaeota*<br>*Thermoproteota*<br>*Unclassified* |
| | **OW** | 1 of 3 phyla | 0.300 | *Euryarchaeota* |
| | **RL** | 1 of 3 phyla | 0.031 | *Euryarchaeota* |

**Table S2. Number of ASVs and observations for bacterial (32), micro-eukaryotic (38) and archaeal (3) phyla included in the Gradient Forest analysis.**

| Bacteria | ASVs | Obs. | Micro-Eukarya | ASVs | Obs. | Archaea | ASVs | Obs. |
|---|---|---|---|---|---|---|---|---|
| *Acidobacteria* | 404 | 215168 | *Annelida* | 1 | 81 | *Euryarchaeota* | 181 | 2043062 |
| *Actinobacteria* | 2461 | 1919765 | *Arthropoda* | 12 | 218 | *Thermoproteota* | 185 | 2975583 |
| *Armatimonadetes* | 39 | 2644 | *Ascomycota* | 59 | 11474 | *Unclassified* | 1 | 25 |
| *Bacteroidetes* | 1727 | 1074007 | *Basidiomycota* | 59 | 9439 | | | |
| *BRC1* | 47 | 19302 | *Blastidiomycota* | 2 | 293 | | | |
| *Chloroflexi* | 1054 | 531049 | *Cercozoa* | 84 | 144871 | | | |
| *Cyanobacteria* | 226 | 70833 | *Chlorophyta* | 72 | 490600 | | | |
| *Deinococcus-Thermus* | 90 | 54346 | *Choanoflagellida* | 2 | 652 | | | |
| *Dependentiae* | 67 | 2574 | *Chytridiomycota* | 24 | 6434 | | | |
| *Elusimicrobia* | 77 | 447 | *Ciliophora* | 27 | 119674 | | | |
| *Entotheonellaeota* | 2 | 961 | *Cryptomonadales* | 1 | 334 | | | |
| *Epsilonbacteraeota* | 1 | 5 | *Cryptomycota* | 2 | 165 | | | |
| *FBP* | 142 | 16286 | *Dinoflagellata* | 21 | 28621 | | | |
| *Fibrobacteres* | 16 | 976 | *Discosea* | 1 | 211 | | | |
| *Firmicutes* | 541 | 15975 | *Ichthyosporea* | 1 | 973 | | | |
| *Gemmatimonadetes* | 563 | 430463 | *Incertae_Sedis* | 2 | 142 | | | |
| *Halanaerobiaeota* | 2 | 50 | *Klebsormidiophyceae* | 1 | 32 | | | |
| *Hydrogenedentes* | 9 | 227 | *Labyrinthulomycetes* | 1 | 12258 | | | |
| *Kiritimatiellaeota* | 1 | 140 | *MAST-12* | 1 | 143 | | | |
| *Lentisphaerae* | 1 | 38 | *Mollusca* | 1 | 19 | | | |
| *Nitrospirae* | 21 | 10072 | *Mucoromycota* | 9 | 541 | | | |
| *Omnitrophicaeota* | 2 | 35 | *Nematoda* | 5 | 10796 | | | |
| *Patescibacteria* | 326 | 28144 | *Nucleariidae & Fonticula Group* | 7 | 1100 | | | |
| *Planctomycetes* | 848 | 134966 | *Ochrophyta* | 116 | 557674 | | | |
| *Proteobacteria* | 3393 | 856725 | *Opisthokonta* | 2 | 16006 | | | |
| *Rokubacteria* | 2 | 34 | *Pavlovophyceae* | 3 | 9262 | | | |
| *Tenericutes* | 13 | 172 | *Peronosporomycetes* | 17 | 6712 | | | |
| *Verrucomicrobia* | 456 | 54344 | *Phragmoplastophyta* | 30 | 14293 | | | |
| *WPS-2 (Eremiobactereota)* | 7 | 720 | *Protalveolata* | 3 | 17021 | | | |
| *WS2* | 1 | 20 | *Prymnesiophyceae* | 1 | 673 | | | |
| *Unclassified* | 184 | 17171 | *Rotifera* | 4 | 20102 | | | |
| | | | *Schizoplasmodiida* | 2 | 79 | | | |
| | | | *Tardigrada* | 2 | 112 | | | |
| | | | *Tubulinea* | 4 | 1037 | | | |
| | | | *Unclassified* | 920 | 2314321 | | | |
| | | | *Vertebrata* | 6 | 4000 | | | |
| | | | *Zoopagomycota* | 1 | 40 | | | |
